# Auditory cortical feedback signals are modulated during songbird courtship

**DOI:** 10.1101/2023.08.08.552485

**Authors:** Caleb Jones, Jesse H. Goldberg

## Abstract

Auditory feedback is important for vocal learning and control, but it remains unclear how the presence of an audience affects neural representations of self-produced sounds. Here we recorded neural activity from the auditory pallium in zebra finches practicing singing alone and directing courtship songs to females. We first discovered that many auditory neurons changed their singing-related discharge patterns during courtship singing. We used syllable-targeted distorted auditory feedback (DAF) to test how signals related to song feedback distortion depend on courtship context. Though past work showed that dopamine neurons uniformly reduce DAF responses during courtship, pallial auditory neurons exhibited heterogeneous DAF signal re-tuning in the presence of the female. Thus, single neurons in the songbird auditory pallium process feedback from self-produced actions differently when singing alone compared to when singing female-directed courtship song.

## Introduction

Motivational drive affects attention and associated sensory responses to external events (Allen et al., 2019; Aton, 2013; Hindmarsh et al., 2021). Hungry or thirsty animals are more alert to food or water cues, which evoke larger brainwide neural responses (Allen et al., 2019; Burgess et al., 2018). Courtship also influences behavioral and neural responses to sensory events (Sakata and Brainard, 2009, 2008; Skals et al., 2005; Zhang et al., 2018), but it remains unclear how representations of self-produced behaviors change with the presence of an audience during courtship. Here we recorded auditory cortical neurons in birds singing alone and to females to test how auditory feedback signals depend on courtship-associated changes in motivational state.

Adult male zebra finches practice undirected song alone and direct courtship song to females (Kao et al., 2008; Zann, 1996). Both undirected and directed songs combine introductory notes, calls, and a stereotyped sequence of phonologically distinct syllables called motifs (Hyland and Tchernichovski, 2019; Rajan and Doupe, 2013). The acoustic structure of undirected and directed motifs is highly similar in adult finches, enabling singing-related neural activity to be precisely aligned and compared across contexts (Kao et al., 2008; Woolley et al., 2014; Zann, 1996). The first goal of this study was to examine motif-aligned discharge in auditory cortical neurons to test if neural representations depend on courtship state.

A second goal was to test how song distortions are represented across performance modes. In past work, distorted auditory feedback (DAF) targeted to specific syllables has been used to drive syllable-specifc vocal plasticity (Andalman and Fee, 2009; Tumer and Brainard, 2007). Though DAF does not capture all aspects of natural song learning, it brings song evaluation under experimental control (Ali et al., 2013; Canopoli et al., 2014; Hoffmann et al., 2016). Birds modify their song to avoid DAF, and several studies support a reinforcement learning framework. Dopaminergic (DA) projections from the ventral tegmental area to Area X, the striatal nucleus of the song system, are necessary for both natural and DAF-based learning (Duffy et al., 2022; Gadagkar et al., 2016; Hisey et al., 2018; Hoffmann et al., 2016; Xiao et al., 2018a; Ali et al., 2013; Harding, 2004; Scharff and Nottebohm, 1991).

Recently, we discovered that DA signals retune during courtship. Specifically, DAF-associated DA signals known to evaluate undirected song are uniformly reduced during female-directed singing - as if a bird does not attend to his own mistakes (Roeser et al., 2023). This discovery raises two possibilities. First, DAF-related DA signals might be retuned at the level of VTA, for example by behavioral state-dependent modulation of intrinsic or synaptic excitability of VTA DA neurons (Xiao et al., 2018b). If this is the case, then DAF responses in upstream inputs to VTA could exhibit similar tuning to auditory feedback during directed and undirected song, but their influence on DA spiking is altered. Another possibility is that auditory processing differs when alone versus when courting a female, analogous to how motivational drives such as thirst or hunger retune widespread neural responses to water or food cues (Allen et al., 2019; Burgess et al., 2018), or how auditory cortical signaling is acutely affected by attention (Ahissar et al., 1992; Hubel et al., 1959) and context (Fritz et al., 2003; Jaramillo et al., 2014). This idea predicts that auditory representations even in a primary pallial area would change with possible attentional and motivational changes associated with a transition from singing alone to singing to a potential mate.

To test these possibilities, we recorded single neuronal activity in the auditory pallium of birds singing alone and to females. We targeted our recording electrodes to Field L, a primary auditory pallial area that projects into multiple higher auditory areas that, in turn, project to VTA (Bottjer et al., 2000; Foster and Bottjer, 1998; Mandelblat-Cerf et al., 2014; Moore and Woolley, 2019). Field L is composed of multiple subdivisions and surrounds the interfacial nucleus and because the implanted wire bundles spread in a small radius of up to ∼0.5 mm, our recordings likely included large territories of the auditory pallium (Figure S1). Although Field L is classically described as a primary auditory thalamorecipient region, its activity may also be shaped by contextual signals related to the courtship context, potentially via recurrent interactions with higher-order auditory forebrain regions such as the caudal mesopallium (CM) and caudomedial nidopallium (NCM) (Bauer et al., 2008; Figure S1C).

Here we report that auditory responses to both bird’s own song and DAF changed in heterogeneous ways at the transition from undirected to courtship directed singing. Thus, a bird’s auditory representations of its own song vary across practice and courtship performance modes. Together with past work showing a re-tuning of premotor signals during courtship (Kao et al., 2008; Singh Alvarado et al., 2021; Woolley et al., 2014; Hessler and Doupe, 1999), our discovery that a sensory area retunes is consistent with distributed changes in sensorimotor processing during courtship.

## Results

### Singing-related auditory discharge is altered during courtship

Because we were interested in how auditory activity changed during female directed song, we recorded at least 20 song motifs during undirected singing and then presented females to elicit courtship song (n=147 neurons, n=11 birds). Consistent with past work (Keller and Hahnloser, 2009), we observed heterogenous singing-related firing in pallial auditory neurons, ranging from highly temporally precise to variable discharge across motif renditions (Figure 1A-C). To compare neural responses between undirected and directed singing, we focused on motif-aligned activity to compare acoustically similar vocalizations in the undirected and directed states (Figure 1A-F). For each neuron, we analyzed how mean firing rates, burst fraction, and the temporal precision of motif-locked discharge depended on courtship state, using analyses previously described (Kao et al., 2008; Sakata and Brainard, 2009; Woolley et al., 2014). Burst fraction was computed as the fraction of spikes occurring in burst events, defined as three (or more) spikes with two (or more) consecutive interspike intervals less than the 25th percentile of the ISI distribution. The temporal precision of neuronal song-locked firing was computed as the intermotif correlation coefficient (IMCC) using a 20 ms Gaussian kernel for smoothing (Goldberg and Fee, 2010; Olveczky et al., 2005; Kao et al., 2008; Sakata and Brainard, 2009; Woolley et al., 2014).

**Figure 1.**
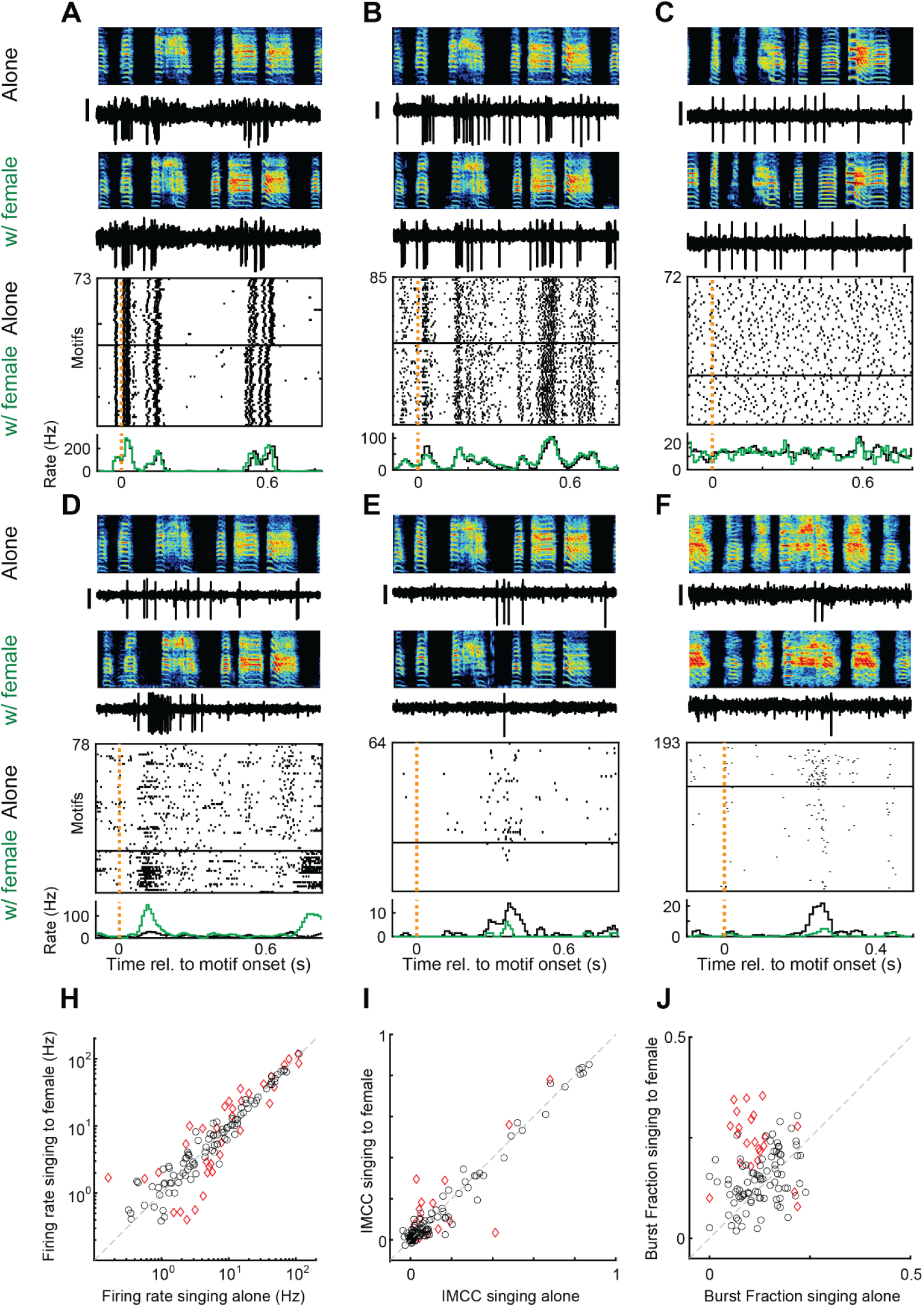
Singing related discharge can retune during courtship. (A-C) Three example neurons with stable firing properties across alone and female-directed song conditions. Top to bottom: single trial example spectrograms, spike discharge, corresponding spike raster plots, and rate histograms (aligned to motif onset) from motif renditions singing alone (black) and to the female (green). Black vertical scale bar for spiking activity is 0.3 mV, y-axis limits of spectrograms are 0.2 to 8 kHz. (D-F) Data plotted as in A-C for three example neurons with significant context-dependent changes in firing. (H-J) Scatter plots of mean firing rates (H), IMCC values (I) and neuronal burst fraction (J) for 138 neurons when singing alone and to the female. Red diamonds: neurons with significant change across conditions (p<0.01, shuffled permutations).

At the population level, neither the temporal precision nor mean firing rate exhibited significant shifts in one direction (Figure 1H-I; Paired t-test, FR: p=0.14, IMCC: p=0.99, n=138 neurons). A small but significant increase in burst fraction was observed (paired t-test, p=9.3x10-6, n=138 neurons, mean ± SEM: 0.11 ± 0.006 vs 0.15 ± 0.007, Figure 1J). Thus auditory neurons did not uniformly change their singing-related firing patterns at the transition to courtship singing, but individual neurons could exhibit significant changes in discharge with changes in courtship state, including increases or decreases in their average motif-locked firing rate (n=19 increase; n=16 decrease, p<0.01), burst fraction (n=20 increase; n=2 decrease, p<0.01), or precision of timing within the motif (n=9 increase; n=5 decrease, p<0.01) (Figure 1D-F, p values derived from Monte Carlo shuffles by condition).

### Singing-related feedback signals can retune during courtship

Past work showed that Field L neurons can exhibit responses to distorted auditory feedback (DAF) during undirected singing (Keller and Hahnloser, 2009). To test if DAF responses exist and/or retune during courtship, we recorded pallial auditory neurons during lone and directed singing as we delivered syllable-targeted DAF (Andalman and Fee, 2009; Tumer and Brainard, 2007). For each bird, a specific syllable was probabilistically targeted with a 50 ms broadband sound played through speakers surrounding the bird (Chen et al., 2020; Gadagkar et al., 2016; Hamaguchi et al., 2014). To quantify DAF responses across the population of neurons and across conditions, we compared neural activity between randomly interleaved renditions of distorted and undistorted songs. DAF responsiveness within each behavioral condition was assessed using circular-shift permutation tests with max-statistic correction across bins. To test whether the DAF response differed between behavioral contexts, we separately permuted context labels (undirected vs directed) within undistorted and distorted trials, recomputed the bin-wise difference-of-differences contrast, and used the maximum absolute shuffled value across bins to obtain family-wise-error-corrected p-values.

Overall, responses to DAF were highly heterogeneous and variable in magnitude. While, as expected, some Field L neurons robustly responded to DAF with activations following song distortion (Figure 2A,D), some neurons’ responses were more subtle (Figure 2C), suggesting a continuum in DAF responsiveness. Interestingly, neural responses to DAF could be modulated by the courtship context in heterogeneous ways. The neuron in Figure 2.2A was DAF responsive during directed singing, but the magnitude of the response was significantly decreased relative to the DAF response while singing alone.

**Figure 2.**
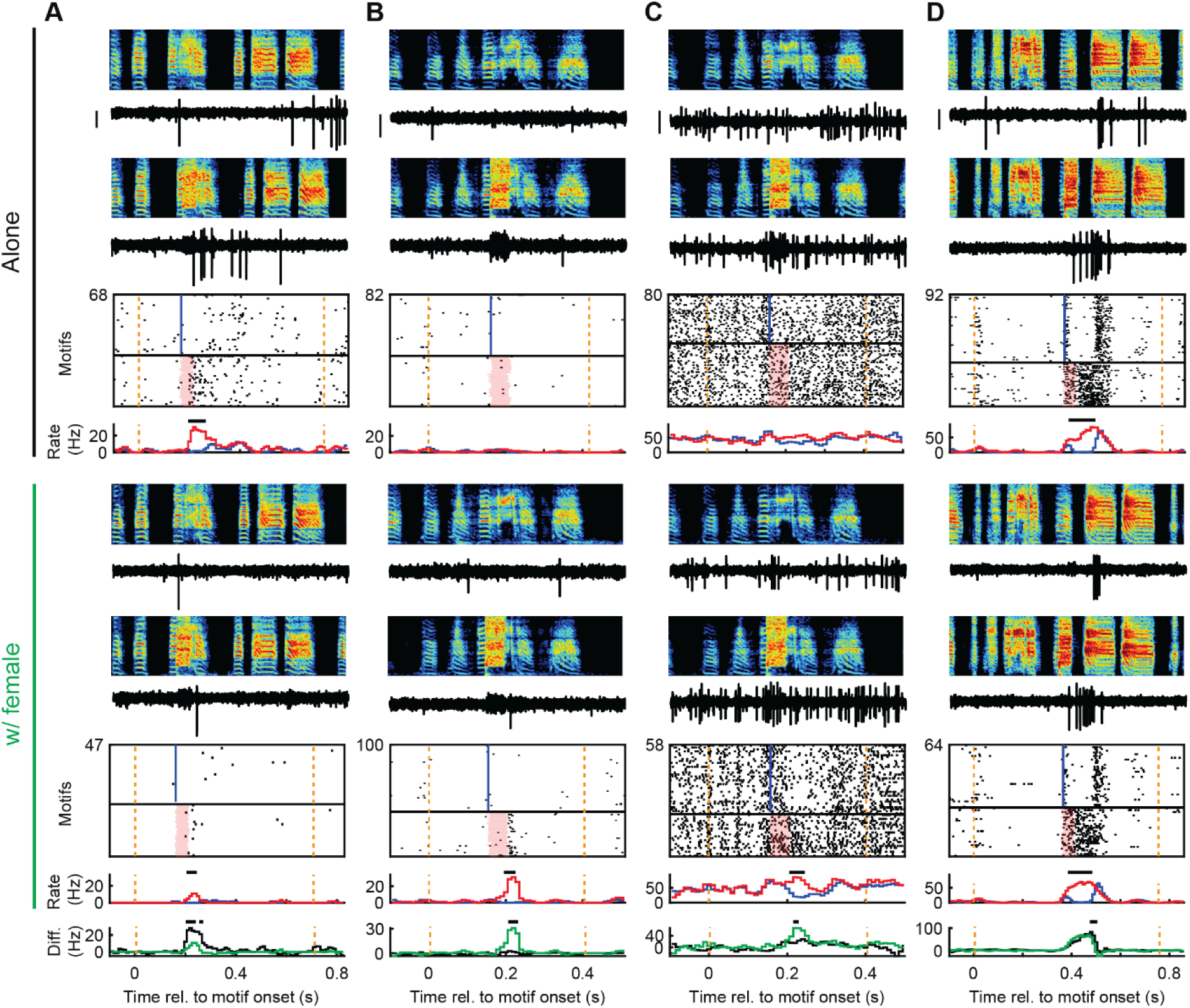
DAF-activated neurons can retune during courtship. (A-D) Four example neurons with DAF responses that vary in the degree to which they are affected by the courtship context. Top to bottom: single trial example spectrograms and spiking activity for undistorted and distorted trials, corresponding raster plots (blue vertical bar denotes feedback target time in undistorted renditions; orange vertical dotted lines denotes onset and offset of song motif). Corresponding rate histograms for undistorted renditions (blue trace) and distorted renditions (red trace). Below are the same plots, but for songs directed to the female. Bottom: difference between undistorted and distorted rate histograms for singing alone (black) and singing to the female (green). All data are time-aligned to motif onset. Black lines above rate histograms and rate difference traces denote bins for significant DAF responses and significant changes in DAF responses, respectively. Scale bar for spiking activity is 0.3 mV. Vertical axis limits for spectrograms are 0.2 to 8 kHz.

Unexpectedly, some neurons (as in Figure 2B) were not DAF responsive during undirected singing, but became DAF responsive during directed singing. The neuron in Figure 2C was moderately DAF responsive during undirected singing, but more so during directed singing. Some neurons, however, as in Figure 2D, exhibited largely unchanged DAF responsiveness across the two social contexts.

Unexpectedly, some neurons were not activated by the song distortion but rather by the lack of target syllable distortion. These activations following undistorted renditions could also depend on the courtship context, as was the case for the neurons in Figure 3A-C, while some neurons were relatively insensitive to the courtship context (Figure 3D).

**Figure 3.**
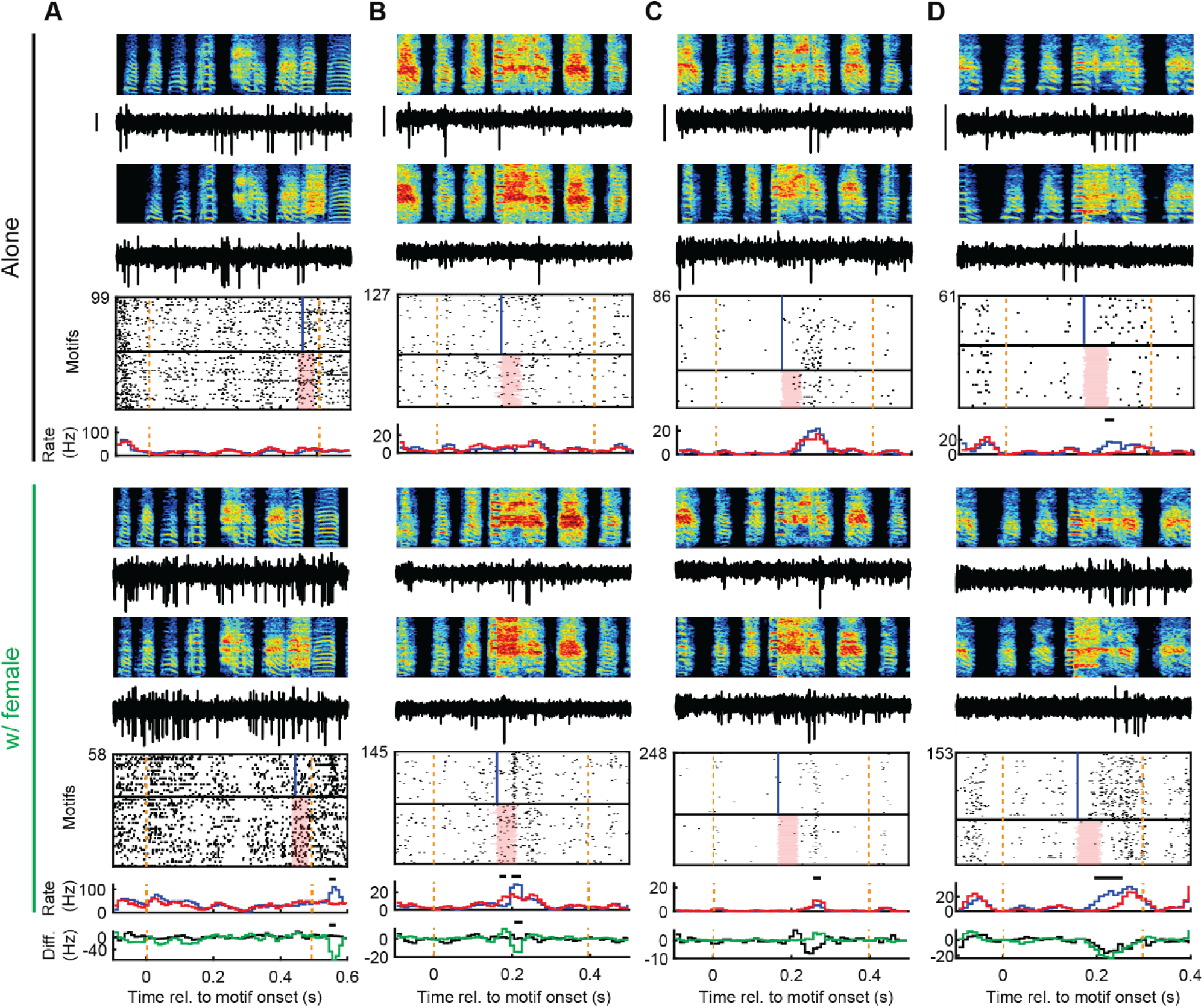
Neurons activated by undistorted singing could depend on courtship context. (A-D) Four example neurons with activations following undistorted renditions that vary in the degree to which they are affected by the courtship context. Top to bottom: single trial example spectrograms and spiking activity for undistorted and distorted trials, corresponding raster plots (blue vertical bar denotes feedback target time in undistorted renditions; orange vertical dotted lines denotes onset and offset of song motif). Corresponding rate histograms for undistorted renditions (blue trace) and distorted renditions (red trace). Below are the same plots, but for songs directed to the female. Bottom: difference between undistorted and distorted rate histograms for singing alone (black) and singing to the female (green). All data are time-aligned to motif onset. Black lines above rate histograms and rate difference traces denote bins for significant DAF responses and significant changes in DAF responses, respectively. Scale bar for spiking activity is 0.3 mV. Vertical axis limits for spectrograms are 0.2 to 8 kHz.

Together, these neural responses exemplify the heterogeneity of DAF-related signaling in the auditory cortex, and that DAF-related signaling can be dependent on the social context in which the bird is performing the song. Importantly, because these neural recordings were performed over long time courses (∼2-8 hours), a strict threshold for stability was imposed. A Pearson’s correlation coefficient of at least 0.99 between the average neural waveform during undirected and directed singing was required for a unit to be considered stable (Dickey et al., 2009; Figure S2). Statistical tests defining auditory neurons as DAF-responsive or not in a binary fashion may not be suitable if the underlying population of DAF-related responses exist on a continuum from responsive to non-responsive. However, to provide a summary of the observed responses, we performed three different analyses. First, we replicated the same analysis as in a previous Field L recording study (Keller and Hahnloser, 2009). This analysis identified 71/147 neurons as DAF responsive in at least one behavioral condition, whereas 76/147 were not responsive in either condition. Next, to visualize changes in DAF responses as a population, we scored each neuron’s DAF response by computing the maximum z-scored difference between undistorted and distorted firing rates in a time window 0-100 ms after distortion target time onset (Figure S3). We found explaining a neuron’s DAF response and change in DAF response across contexts difficult to describe with a single number that was highly sensitive to parameter choices. Because the previously used tests were highly sensitive and did not explicitly test for a change in DAF responsiveness, we adopted the previously mentioned permutation-based test. This more conservative approach identified 48/147 neurons as DAF responsive in at least one condition, with 13 of those neurons exhibiting a significant modulation in their DAF response between undirected and female-directed singing. Together, these data show that auditory responses to distorted auditory feedback during singing can retune during courtship.

A previous study recording from Field L in zebra finches reported neural activations prior to the onset of singing, consistent with premotor signaling (Keller and Hahnloser, 2009). We tested for context dependent changes in premotor activity by examining peaks in neural activity aligned to motif onsets. Across the population neurons did not exhibit significant changes in the timing of motif onset-aligned activity (Figure S4).

Action potential width of individual neurons has previously been used to classify putative interneurons or putative principal cells in the zebra finch auditory pallium (Calabrese and Woolley, 2015). We tested if DAF-response scores, mean firing rates during singing, burst fraction, IMCC, and the change in all of these between undirected and directed singing was correlated with spike half width (spike half-width measured as peak to trough time; Figure S5). DAF-response in either condition, the change in DAF response across conditions, and the change in firing rate, burst fraction, and IMCC were not significantly correlated with spike width (Figure S5A-B). Consistent with previous literature, firing rate was significantly correlated with spike width (Pearson’s correlation, p=0.003; Figure S5C). Interestingly, burst fraction (p=8.3x10-4) and IMCC (p=5.4x10-5) were also significantly correlated with spike width (Figure S5C).

## Discussion

By recording from the auditory pallium in zebra finches singing alone and to females, we discovered, for the first time to our knowledge, that auditory representations of an animal’s own vocalizations change with an audience. These findings extend past work in finches showing that social context affects auditory responses to the calls of conspecifics (Angeloni and Geffen, 2018; Menardy et al., 2014; Remage-Healey et al., 2010). More broadly, these findings support the idea that auditory cortical activity is modulated not just by acoustic features (Gervain and Geffen, 2019; King et al., 2018) but also diverse phenomena such as attention (Hubel et al., 1959), primary rewards (David et al., 2012), task parameters (Downer et al., 2015; Fritz et al., 2003); (Angeloni and Geffen, 2018; Menardy et al., 2014, 2012; Remage-Healey et al., 2010), and even hormone levels (Angeloni and Geffen, 2018; Menardy et al., 2014, 2012; Remage-Healey et al., 2010).

Detecting differences between predicted and actual sensory feedback is a crucial aspect of motor learning (Keller and Mrsic-Flogel, 2018), and auditory cortical neurons in diverse species can signal mismatch errors (Eliades and Wang, 2008; Keller and Hahnloser 2009; Parras et al., 2021; Ulanovsky et al., 2003). However, it is important to note that DAF-related changes in firing in auditory neurons do not necessarily imply that neurons compute sensory prediction errors. DAF-related responses could arise from ordinary auditory tuning to the broadband distortion stimulus. In past recordings from VTA we observed a bimodal distribution of error responses, and the error-responding neurons were the ones that projected to Area X (Gadagkar et al., 2016). Yet in cortex there is uncertainty about the suitability of classifying single neurons by response profile, as task-relevant parameters that can be decoded from neuronal populations may not be apparent when examining single neuronal representations (Kaufman et al., 2014; Mante et al., 2013; Rigotti et al., 2013; Williams et al., 2018). We observed a continuum of DAF responses during both directed and undirected singing which made it difficult to unambiguously define a neuron as being DAF responsive or not.

Because the main goal of this study was to test if courtship-associated reduction in DAF signaling, recently observed in VTA DA neurons (Roeser et al., 2023), resulted from a local process in VTA or reflected a retuning of auditory responsiveness, we explicitly tested for changes in DAF responsiveness between alone and female-directed singing. Surprisingly, we discovered that Field L neurons could retune at the transition from lone to courtship singing in diverse ways, consistent with distributed modulation of auditory processing that does not fully explain the uniform DAF-signal attenuation observed in VTA. Interestingly, Area X and LMAN neurons uniformly increase their temporal precision and reduce their burstiness during courtship singing (Kao et al., 2008; Roeser et al., 2023; Sakata and Brainard, 2008; Singh et al., 2021; Woolley et al., 2014). In contrast, auditory pallium neurons could exhibit increases or decreases in burstiness, temporal precision, or mean rate with courtship.

An open question is how auditory pallium receives information about whether a female is present, and how this information influences neural activity. Several neuromodulatory systems with possible information about courtship state project to songbird auditory forebrain, including acetylcholine (Shea and Margoliash, 2010), serotonin (Yip et al., 2020), and norepinephrine (Cardin and Schmidt, 2004). Estrogens can also rapidly modulate auditory firing (Remage-Healey et al., 2010; Scarpa et al., 2022). Yet how the courtship state may be orchestrated across multiple brain regions remains unclear. One possibility is that hypothalamic nuclei such as the medial preoptic nucleus (MPOA) initiate the courtship state. The MPOA projects to multiple brainstem neuromodulatory areas which in turn project broadly throughout the forebrain, including the song system and auditory system (Riters and Alger, 2004; Ben-Tov et al., 2023; Castelino and Ball, 2005; Singh Alvarado et al., 2021). It will be interesting in future studies to examine the neural signals propagating through these pathways at transitions into and out of the courtship state.

## Materials and Methods

### Subjects

Sixteen adult male zebra finches (>90 days post hatch) were the subjects of this study. Animal care and experiments were carried out in accordance with NIH guidelines and were approved by the Cornell Institutional Animal Care and Use Committee.

### Surgery and awake-behaving electrophysiology

For chronic neural recordings, subjects were anesthetized with isoflurane inhalation and mounted on a stereotaxic instrument for probe implantation. All probes were 16-channel moveable electrode bundles (Innovative Neurophysiology). Probes were implanted 1.5-2.0 mm anterior and 1.5-2.0 mm lateral of the bifurcation of the mid-sagittal sinus to target Field L (Keller and Hahnloser 2009) at a head angle of 80 degrees (measured as the angle from the tip of the beak to the center of the ear bars relative to the horizontal plane). The end of the cannula was implanted 1.5 mm ventral to the surface of the brain. Birds were then placed alone in a sound isolation chamber (12 hour light/dark cycle) with ad libitum food and water and allowed 1 day to recover post op before being subjected to the distorted auditory feedback (DAF) protocol. 2-3 days were allotted for habituation to DAF and to ensure the bird began to spontaneously sing enough motifs in social isolation before neural recordings. Electrode bundles were extended from the end of the cannula on a daily basis by ∼50 μm. DAF was implemented with a custom LabView acquisition program that analyzed song syllables in real-time and delivered syllable-targeted feedback. DAF (50 ms broadband noise bandpass filtered at 1.5-8 kHz to match frequency range of zebra finch song) was played over speakers in the recording chamber on top of a specific target syllable randomly on 50% of motif renditions. Experiments were carried out in the male’s home cage, which was inside a sound isolation chamber. When the homecage lights came on each day, recording began and the male was left alone to sing at least 40 undirected song motifs. Female directed motifs were then recorded by presenting a female in the chamber in ∼10 minute intervals throughout the day until at least 40 directed song motifs were elicited (Roeser et al., 2023). Electrode placement was verified at the end of the experiment with small electrolytic lesions, histology, and dark field imaging. 11 of the 16 implanted birds yielded single unit recordings and sang sufficient motifs for the experiment. Many channels on the probes recorded multi-unit activity, which were taken note of but not analyzed in this study.

### Neural recording and analysis

Neural signals were acquired with the Intan RHD recording controller and 16-channel Intan headstages that directly interfaced with the moveable bundles. Sampling rate was set to 20kHz and recording was manually controlled with the Intan recording software, where a 60 Hz notch filter was applied and spiking activity of single units could be visualized in real time. Audio data and a real-time digital copy of the DAF signal were simultaneously recorded with neural data in the recording controller such that all data could be easily time aligned. A custom MATLAB GUI was used for visualizing song and neural data, and for spike sorting. Neural recordings were bandpass filtered between 0.4 kHz and 6-8 kHz and single unit spiking activity was manually sorted as previously described (Goldberg and Fee, 2010). Initially, 161 neurons were recorded, but only highly stable neurons with a correlation coefficient of at least 0.99 between the average waveform during undirected and directed singing were kept (Dickey et al., 2009). Spike half width was measured as peak to trough time on the average waveform.

Motif aligned spiking activity was time-warped to the median duration of undirected or directed motifs. Any motifs that had overlapping time with a female call in directed motifs was excluded from analysis. Firing rate (FR) histograms were computed by binning spiking events in 10 ms windows and smoothing with a 3-bin moving average. To calculate the significance of the FR changes across behavioral contexts, we randomly assigned spike trains from each song motif trial as undirected or directed groups while conserving trial numbers, then calculated FR values for the randomized dataset, as previously described (Goldberg and Fee, 2010). This shuffling without replacement was repeated 10,000 times for each neuron, yielding 10,000 new changes in FR values. Measured changes in firing rate that were greater than the 99th percentile of the shuffled distribution were considered significant, as in previous studies (Sakata and Brainard, 2008). This procedure was repeated for IMCC and burst fraction. A p value less than 0.01 was considered significant to account for multiple comparisons. To quantify the degree to which neuronal firing was time-locked to song, the intermotif correlation coefficient (IMCC) was calculated as described previously (Chen et al., 2019; Kao et al., 2008; Olveczky et al., 2005). To compute IMCC, motif-aligned FR was mean-subtracted and smoothed with a Gaussian kernel of 20 ms SD, resulting in a rate vector, ***r_i_***, for each motif. IMCC was defined as the mean of all pairwise correlation coefficients between ***r_i_*** as follows:

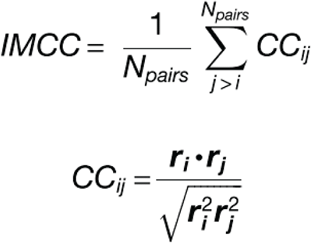

We defined bursts as events containing three or more spikes with consecutive interspike intervals less than the 25th percentile of the 25th percentile of the interspike interval distributions during singing. Burst fraction was calculated as the fraction of spikes during song motifs that occurred during bursts.

We quantified the DAF responsiveness for each neuron in two ways, following approaches previously described (Chen et al., 2019; Gadagkar et al., 2016; Keller and Hahnloser, 2009). Spike counts in 30 ms time windows shifted in 5 ms steps up to 50 ms after DAF offset were generated for undistorted and distorted trials. A WRS test assessed the significance of each 30 ms window, and only neurons with 2 or more consecutively significant (p<0.05) windows were considered a significant DAF response (Keller and Hahnloser, 2009).

To more stringently test for DAF responsiveness and explicitly test for changes in DAF responsiveness between undirected and directed singing, we used the motif-aligned, time-warped spike trains counted in 10ms bins, smoothed over 3 bins, and defined bin-wise DAF-response vectors as distorted minus undistorted activity within the undirected and directed conditions. A bin-wise difference-of-differences vector was used to quantify context-dependent changes in the DAF response. Within-condition DAF responsiveness was tested using circular-shift permutation tests in which each trial’s spike-count vector was independently circularly shifted by a random number of bins, preserving within-trial temporal structure while disrupting stimulus alignment (Harrison and Geman, 2011; Stella et al., 2022; Amarasingham et al., 2011). For each shuffle, shuffled DAF-response vectors were recomputed and the maximum absolute value across bins was stored to generate max-statistic null distributions. To test whether DAF responsiveness differed between contexts, context labels were randomly permuted separately within undistorted and distorted trials, shuffled DAF-response vectors were recomputed for each context, and the shuffled difference-of-differences vector was calculated; again, the maximum absolute value across bins was stored on each shuffle. Family-wise-error-corrected bin-wise p-values were obtained by comparing the absolute observed statistic at each bin to the corresponding null distribution of shuffled maxima.

To investigate premotor related activity, we aligned spikes to motif onsets that had at least 30 ms of preceding silence. Spikes were binned in 3ms bins and smoothed over 3 bins. Trial-averaged firing rates were computed and peaks were detected using MATLAB’s findpeaks function. The significance of a peak was determined by circularly shuffling the spike trains and recomputing the maxima, yielding a null distribution to compare the measured value against. All peaks with p<0.05 were considered significant, and the earliest peak in firing rate was plotted in Figure S4A.

---

**Figure S1.**
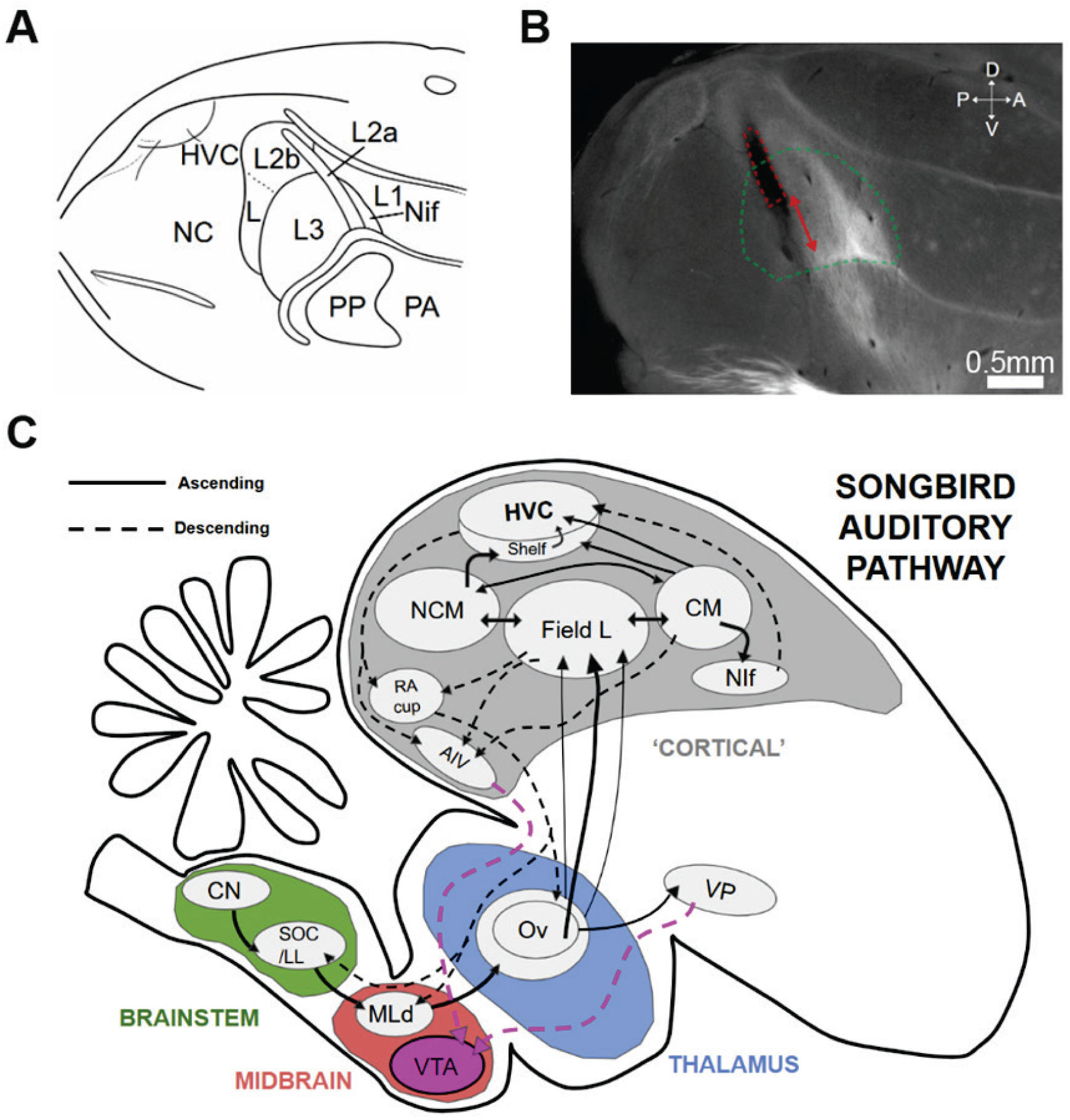
Auditory system of the zebra finch and example histology. (**A**) Example sagittal view of the anatomical structure of auditory areas in the zebra finch brain, adopted from a previous study (Fortune and Margoliash, 1992). (**B**) Example brain slice approximately 1.5mm medial from the midsagittal sinus showing the wire bundle implant location (dotted red line) and the travel path of the moveable bundle (red arrows). Green dotted line denotes the approximate region recorded across all birds, estimated from histology of each bird (**C**) Connectivity diagram showing the ascending and descending connections between auditory areas in the zebra finch brain. Pink lines show connections to the VTA. Acronyms: CN; cochlear nucleus, SOC; superior olivary complex, MLd; dorsal mesencephalic nucleus, VTA; ventral tegmental area, Ov; nucleus ovoidalis, VP; ventral pallidum, CM; caudal mesopallium, NIf; interfacial nucleus, NCM; caudomedial nidopallium, RA; robust nucleus of the arcopallium, AIV; ventral portion of the intermediate arcopallium. Connections between areas adopted from connectivity diagrams in previous studies (Shaevitz and Theunissen, 2007; Vates et al., 1996; Bauer et al., 2008).

**Figure S2.**
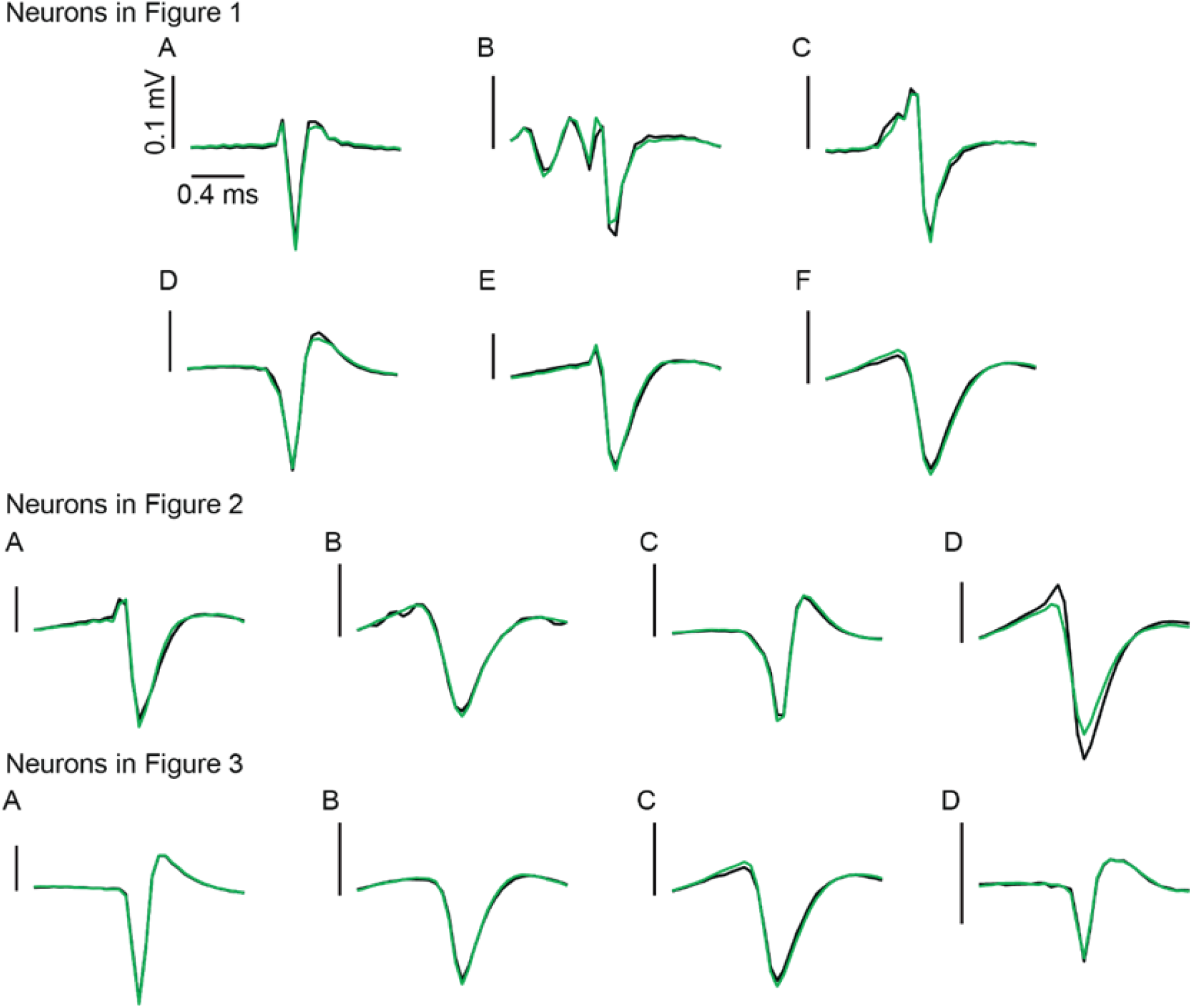
Spike waveforms are stable across undirected and directed singing. Average waveforms for single units from Figures 2.1-2.3 during undirected singing (black) and directed singing (green). All units included in this study had a Pearson’s correlation coefficient between the undirected and directed average waveforms of >0.99.

**Figure S3.**
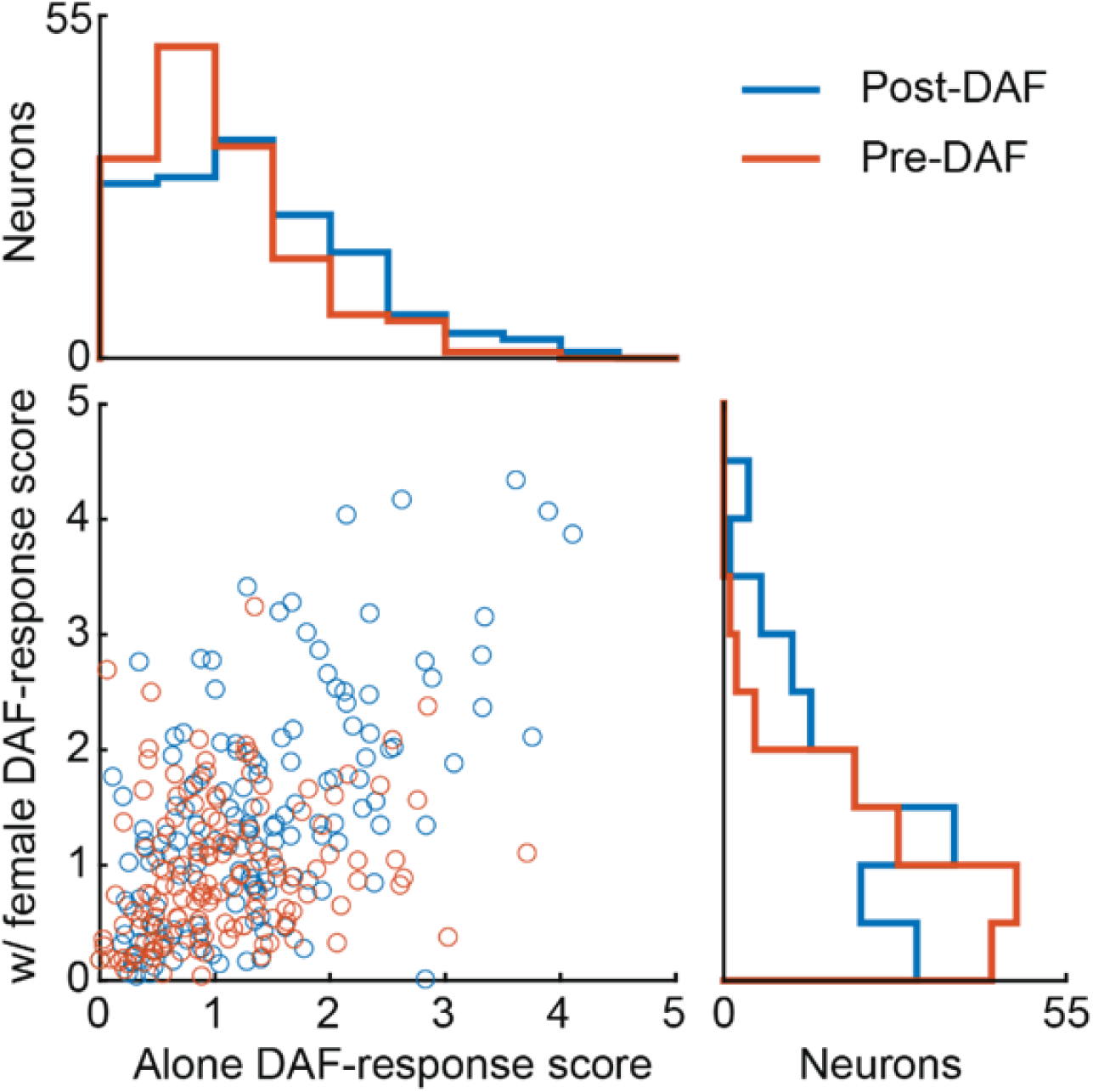
Distributions of absolute DAF-response scores in pallial auditory neurons. Scatter plot where each dot represents the absolute z-scored DAF-response of a single neuron during the 100 ms interval after the onset of DAF (blue) and 100 ms before DAF onset (orange). Corresponding histograms for each condition are projected along the x and y axes.

**Figure S4.**
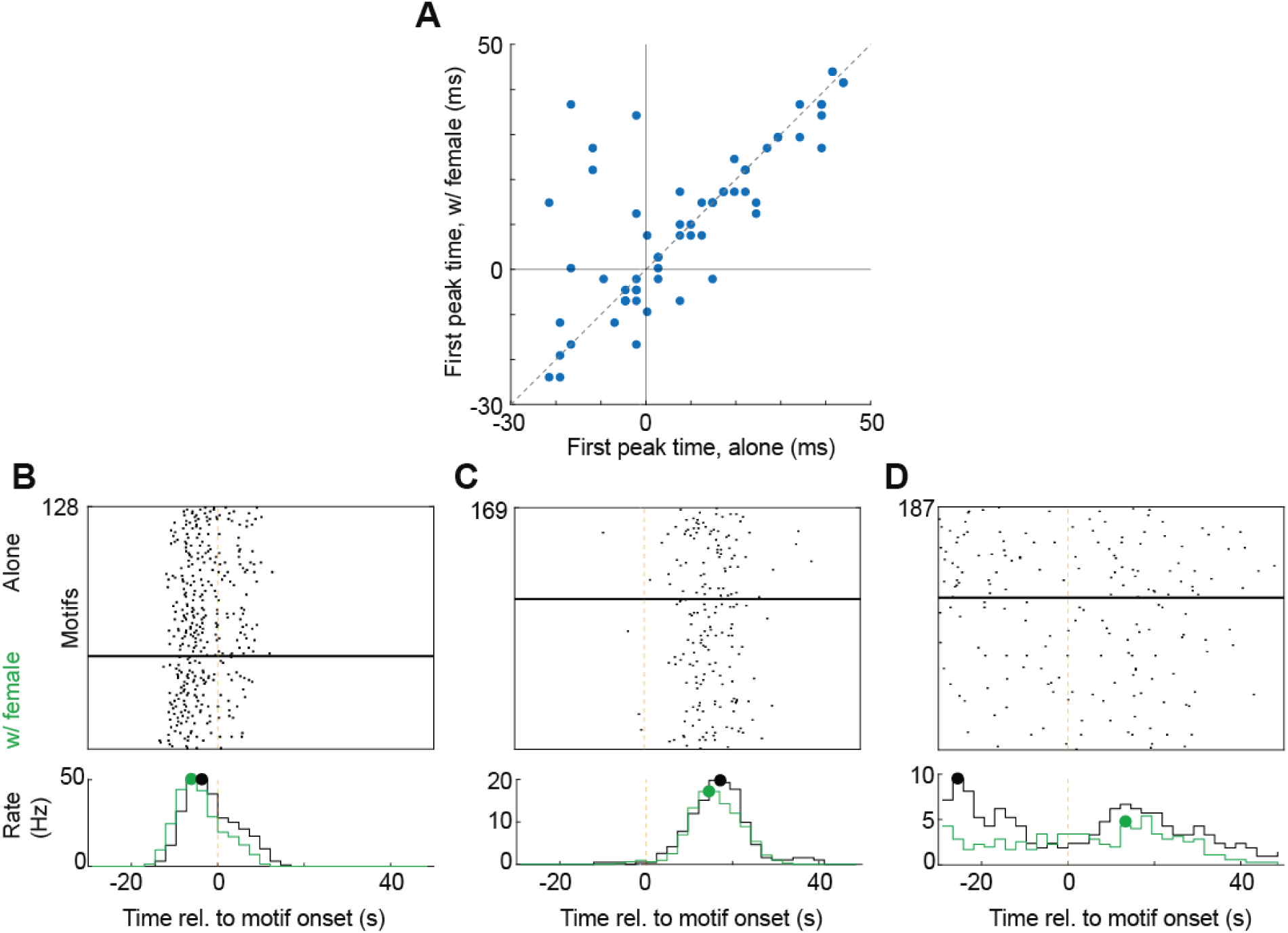
Latencies of neural response vary from pre- to post-motif onset and are largely stable across conditions. (**A**) Scatter plot of latencies of first firing rate peak between undirected and directed singing. (**B**) Example neuron with activity before motif onset; raster of spike times and average firing rates for undirected (black) and directed (green) aligned to motif onset. Green and black dots indicate the earliest significant peak. (**C**) Example neurons plotted as in (B) but for a neuron responding post motif onset. (**D**) Example neuron plotted as in (B-C) but for a neuron with a relatively larger change in peak response time.

**Figure S5.**
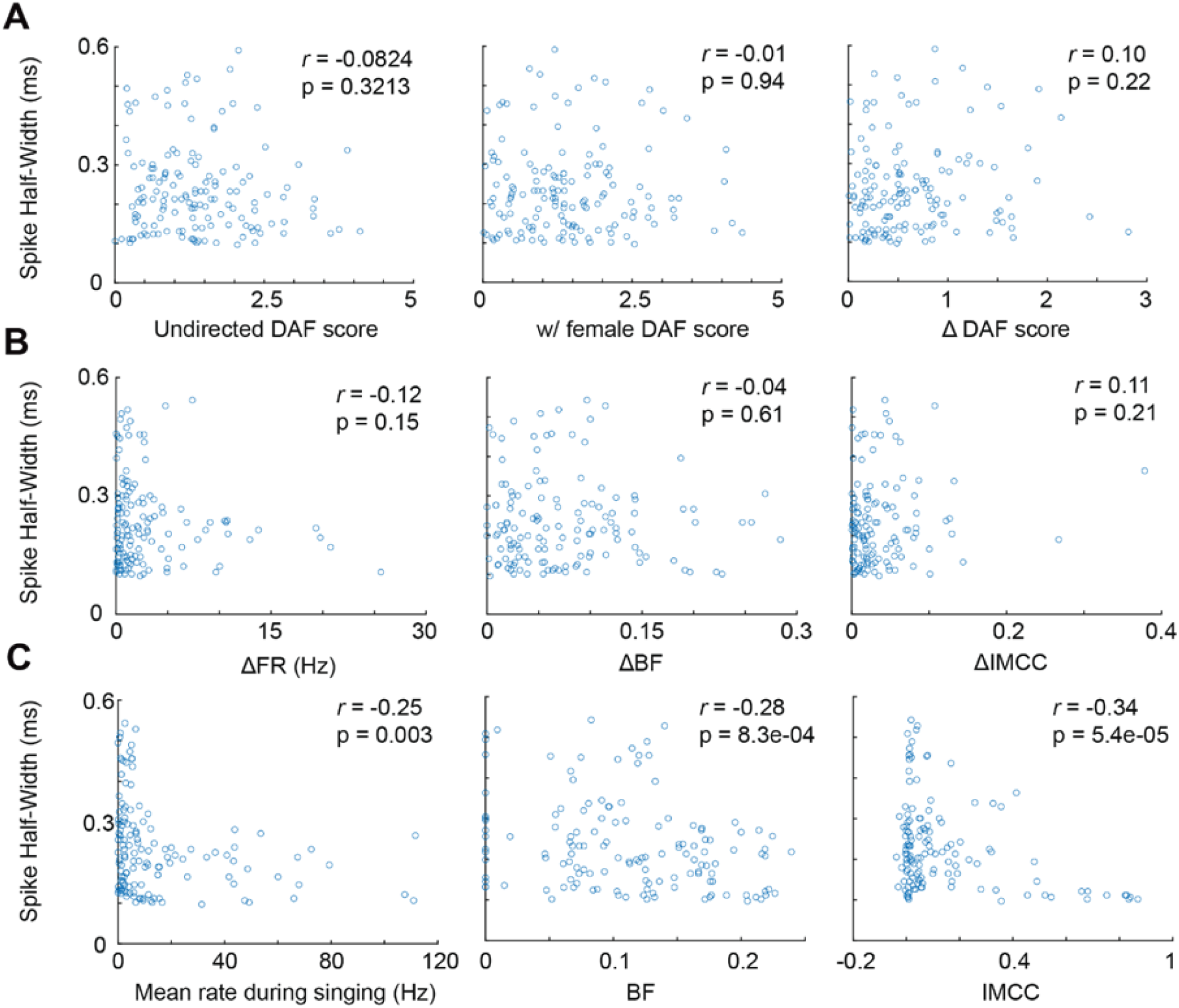
Spike half-width correlations with neural discharge properties. (**A**) Scatter plots of DAF-response scores in undirected and directed, and the change between them (undirected-directed) vs spike half-width with no significant correlations (**B**) Scatter plots of the change in firing rate, burst fraction (BF) and IMCC (undirected-directed) vs spike half-width with no significant correlations. (**C**) Scatter plots of mean firing rate, burst fraction, and IMCC, all during undirected singing, vs spike half-width. All correlations in (C) are significant with correction for multiple comparisons (uncorrected p-values shown). Pearson’s linear correlation coefficients and associated p values are shown in each plot.

